# pathfindR: An R Package for Pathway Enrichment Analysis Utilizing Active Subnetworks

**DOI:** 10.1101/272450

**Authors:** Ege Ulgen, Ozan Ozisik, Osman Ugur Sezerman

**Author notes:** Correspondence to: Ege Ulgen, Address: Department of Biostatistics and Medical Informatics, School of Medicine, Acibadem Mehmet Ali Aydinlar University, Istanbul, Turkey.

## Abstract

**Summary:** PathfindR is a tool for pathway enrichment analysis utilizing active subnetworks. It identifies gene sets that form active subnetworks in a protein-protein interaction network using a list of genes provided by the user. It then performs pathway enrichment analyses on the identified gene sets. Further, using the R package pathview, it maps the user data on the enriched pathways and renders pathway diagrams with the mapped genes. Because many of the enriched pathways are usually biologically related, pathfindR also offers functionality to cluster these pathways and identify representative pathways in the clusters. PathfindR is built as a stand-alone package but it can easily be integrated with other tools, such as differential expression/methylation analysis tools, for building fully automated pipelines. In this article, an overview of pathfindR is provided and an example application on a rheumatoid arthritis dataset is presented and discussed.

**Availability:** The package is freely available under MIT license at: https://github.com/egeulgen/pathfindR

## 1. Introduction

High-throughput technologies have revolutionized biomedical research by enabling comprehensive characterization of biological systems. These technologies allow researchers to identify a list of differentially expressed genes/proteins or differentially methylated genes, which most likely play a role in the formation of the phenotype.

However, this list often falls short of providing mechanistic insights into the underlying biology of the disease being studied^1^. Therefore, we face a challenge posed by high-throughput experiments: extracting relevant information that allows us to understand the underlying mechanisms from a long list of genes or shortly, finding a needle in the haystack.

One approach, which reduces the complexity of analysis while simultaneously providing great explanatory power, is identifying groups of genes that function in the same pathways, i.e. pathway analysis^1^. Pathway analysis has been successfully and repeatedly applied to gene expression^2,3^, proteomics^4^ and DNA methylation data^5^, in addition to various other applications^6^-^10^.

However, there are drawbacks to pathway analysis. Most importantly, the statistics used by pathway analysis approaches usually consider the number of genes in a list alone and are independent of the values associated with genes, such as fold-changes or p values. By treating each gene equally, they also assume that each gene is independent of the other genes. Because they ignore information on interactions of genes, directly performing pathway analysis on a gene set is not completely informative.

For a given list of significant genes, an active subnetwork is defined as a group of interconnected genes in a protein-protein interaction network (PIN) that mostly consists of significant genes. In short, active subnetworks define distinct disease-associated sets of interacting genes. For the identification of active subnetworks, various algorithms have been proposed, such as greedy algorithms^11^-^19^, simulated annealing^20^-^21^, genetic algorithms^22^-^26^ and mathematical programming-based methods^27^-^31^.

With pathfindR, we propose to leverage interaction information from active subnetworks to extract the most relevant pathways, utilizing both the p values of individual genes and information from a PIN. In the pathfindR approach, information from four resources are integrated to determine the mechanisms underlying the disease: (i) differential expression/methylation information obtained through omics analyses, (ii) interaction information through the protein-protein interaction network, (iii) Kyoto Encyclopedia of Genes and Genomes (KEGG)^32,33^ pathways, (iv) clustering of related pathways and establishment of representative pathways.

The pathfindR package was developed based on a previous approach developed by our group for genome-wide association studies (GWASes): Pathway and Network-Oriented GWAS Analysis (PANOGA)^34^. PANOGA was successfully applied to uncover the underlying mechanisms in GWASes of various diseases, such as rheumatoid arthritis^35^, intracranial aneurysm^36^, epilepsy^37^ and Behcet‘s disease^38^. pathfindR applies an approach similar to PANOGA to “omics” experiments with additional functionality, further described below.

In this article, we present the details on pathfindR along with an example application on rheumatoid arthritis (RA) differential expression data.

## 2. Methods

### 2.1. The pathfindR Case Study - Analysis on RA Data

The dataset GSE15573 was obtained from the National Center for Biotechnology Information (NCBI) - Gene Expression Omnibus (GEO). This dataset aimed to characterize gene expression profiles in the peripheral blood mononuclear cells of 18 RA patients versus 15 healthy subjects. We performed differential expression analysis between these two groups using the R^39^ package limma^40^. The differentially-expressed genes (DEGs) with adjusted p values ≤ 0.05 (n = 571) were used to create the example input dataset *RA_input*. This dataset includes the gene symbols, log-fold-change values and adjusted p values for the DEGs.

Active subnetwork search and enrichment analysis of the RA differential-expression data was performed with pathfindR using the greedy active subnetwork search algorithm and the Biogrid PIN.

Next, the enriched pathways were clustered and representative pathways were obtained.

The analysis approach is explained in detail below.

### 2.2. Protein-protein Interaction Networks

The user can choose between the protein-protein interaction data of KEGG, Biogrid^41,42^, GeneMANIA^43^ and InTact^44^.

The KEGG PIN was created by an in-house script using the KEGG pathways on December 31, 2017. The relations among genes were added to the PIN as undirected links, removing any duplicate interactions.

For the GeneMania PIN, only interactions with weights ≥ 0.0006 were kept, allowing only strong interactions.

All PINs were formatted as simple interaction files (SIFs) for use in analyses.

The user can also use a PIN of their choice by supplying the path of the SIF to the wrapper function *run_pathfindR*.

### 2.3. Scoring of Subnetworks

In pathfindR we followed the scoring scheme that was proposed by Ideker et al.^20^. p value of each gene is converted to z-score using Eq. 1 and the score of a subnetwork is calculated using Eq. 2. In Eq. 2, A is the set of genes in the subnetwork and k is its cardinality.

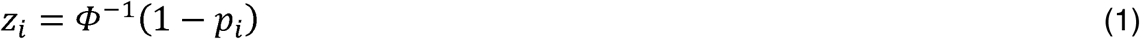

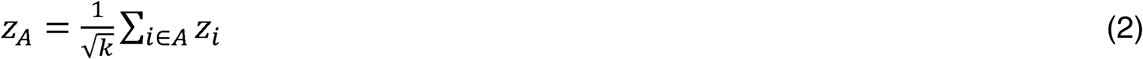

In the same scoring scheme, there is also a Monte Carlo approach for the calibration of the scores of subnetworks against background distribution. Using randomly selected genes, 2000 subnetworks of each possible size are constructed and for each possible size, the mean and standard deviation of the score is calculated. These values are used to calibrate subnetwork score using Eq. 3.

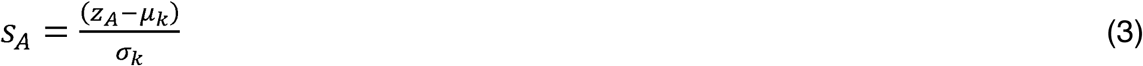

### 2.4. Active Subnetwork Search Algorithms

Currently, there are three algorithms implemented in the pathfindR package for active subnetwork search: greedy algorithm, simulated annealing algorithm and genetic algorithm.

#### 2.4.1. Greedy Algorithm

Greedy algorithm is the problem-solving/optimization concept that chooses locally the best option in each stage with the hope of reaching the global optimum. In active subnetwork search, this is generally applied by starting with a significant seed node and considering addition of a neighbor in each step to maximize the subnetwork score. In pathfindR, we used the approach in Chuang et al.^13^: This algorithm considers addition of a node within a specified distance d to the current subnetwork. In our method maximum depth from the seed can also be set. With the default parameters, our greedy method considers addition of direct neighbors (d=1) and forms a subnetwork with a maximum depth of 1 for each seed. Because the expansion process runs for each significant seed node, several overlapping subnetworks emerge. In pathfindR, overlapping subnetworks are handled by discarding a subnetwork that overlaps with a higher scoring subnetwork more than a threshold, which is set to 0.5 by default.

#### 2.4.2. Simulated Annealing Algorithm

Simulated annealing improves the greedy search by accepting non-optimal actions to increase exploration in the search space. The probability of accepting a non-optimal action decreases in each iteration. In active subnetwork search context, the search begins with a set of randomly chosen genes (that will be referred to as genes in “on” state), connected components in this candidate solution are found and the scores are calculated. In each iteration the state of a random node is changed from on to off, vice versa, connected components are found in the new solution and their scores are calculated. If the score improves, the change is accepted, if the score decreases, the change is accepted with a probability proportional to the temperature parameter that decreases in each step.

#### 2.4.3. Genetic Algorithm

Genetic Algorithm is a bio-inspired algorithm that mimics natural selection by implementing fitness-based parent selection, crossover of genes and mutation. In our genetic algorithm implementation, candidate solutions represent on/off state of each gene. In the algorithm, we used rank selection and uniform crossover. In each iteration, the fittest solution of the previous population is preserved if the highest score of the current population is less than the previous population‘s score. In every ten iterations, the worst scoring 10% of the population is changed with random solutions. Because uniform cross-over and addition of random solutions make adequate contribution to exploration of the search space, mutation is off in default settings.

### 2.5. Active Subnetwork-Oriented Pathway Enrichment Analysis

Our active subnetwork-oriented pathway enrichment is implemented as the wrapper function *run_pathfindR*. The overview of the approach is presented in Figure 1.

**Figure 1:**
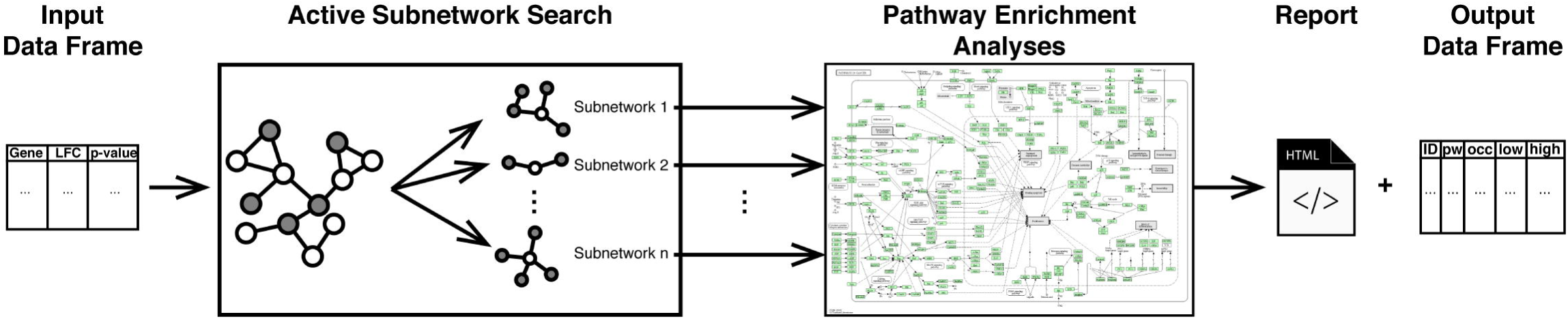
Flow diagram of the pathfindR active subnetwork-oriented pathway enrichment analysis approach

Initially, the input is filtered so that all p values are less than or equal to the given threshold (default is 0.05). Next, gene symbols that are not found in the PIN are identified. If aliases of these gene symbols, obtained through the R package org.Hs.eg.db^45^, are found in the PIN, the symbols are converted to the corresponding aliases.

The processed data is used for active subnetwork search. The identified active subnetworks are then filtered via the following criteria: (i) has a score larger than the given threshold (default is 3) and (ii) contains at least a specified number of DEGs (default is 2).

Using the genes in each of the remaining subnetworks, pathway enrichment analyses are performed via one-sided hypergeometric testing. The enrichment tests use the genes in the PIN as the gene pool. Using the genes in the PIN instead of the whole genome provides more statistical strength because active subnetworks are identified using only the genes in the PIN. The p values obtained from the enrichment tests are adjusted using the Bonferroni method.

Pathways with adjusted p values larger than the given threshold (default is 0.05) are discarded. This process of active subnetwork search and enrichment analysis is repeated for a selected number of iterations (default is 10 iterations for greedy and simulated annealing algorithms, 1 for genetic algorithm). These iterations are executed in parallel via the R package foreach^46^.

Finally, the lowest and the highest adjusted p values, the number of occurrences over all iterations and up-regulated and down-regulated DEGs in each enriched pathway are returned as a data frame. Additionally, Hypertext Markup Language (HTML) format reports with the pathfindR enrichment results, linked to the visualizations of the pathways, as well as the table of converted gene symbols are created. The pathway diagrams are created using the R package pathview^47^. These diagrams display the involved genes colored by change values on a KEGG pathway graph.

### 2.6. Pathway Clustering and Partitioning

Enrichment analysis usually yields a large number of related pathways. In order to establish representative pathways among similar groups of pathways, we propose that clustering can be performed via an approach based on a method described previously by Chen et al^48^. This approach is described below:

Firstly, an overlap index matrix *OI* containing overlap indices between all pairs of pathways is calculated. For each pathway *P*_*i*_ in the dataset, let *G*_*i*_ be the set of all genes in *P*_*i*_. For a pair of pathways *P*_*i*_ and *P*_*j*_, *OI*_*i,j*_ is defined in Eq. 4.

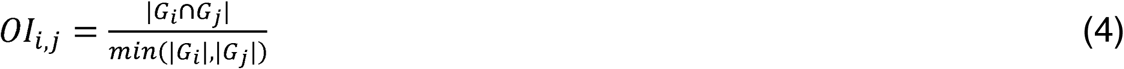

Afterwards, defining each row *o*_*i*_ of the matrix *OI* as the gene overlap profile of pathway

*P*_*i*_, the Pearson correlation coefficients *R*_*i,j*_ are calculated for each pair of *o*_*i*_ and *o*_*j*_. These are then transformed into pairwise distances *PD*_*i,j*_ *= 1 - R*_*i,j*_. This distance calculation approach is implemented in pathfindR as the function *cluster_pathways*. Using this distance metric *PD*, pathways are clustered via hierarchical clustering with the desired agglomeration method. Via a shiny^49^ application, the hierarchical clustering dendrogram is visualized. In this application, the user can select the agglomeration method and the distance value at which to partition the tree. The representative pathway for each cluster is chosen as the pathway with the smallest “lowest p” value. The dendrogram with the cut-off value marked with a red line is dynamically visualized and the resulting cluster assignments of the pathways and annotation of representative pathways are presented as a table. This table can be saved as a comma-separated values (CSV) file.

This clustering and portioning method is implemented as the wrapper function *choose_clusters* in the pathfindR package.

## 3. Results of Analysis on RA Data

In the analysis of the RA differential expression data, pathfindR identified 36 KEGG pathways to be enriched (Table S1). Upon examination of these pathways, some appeared to be biologically related, such as various signaling pathways. Therefore, clustering of these 36 KEGG pathways was performed. Upon manual inspection, the clustering dendrogram was cut at a distance of 0.66 (Figure 2), and 13 representative pathways were obtained (Table 1). Below, we discuss the functional relevance of the identified representative pathways to the pathogenesis of RA.

**Table 1:**
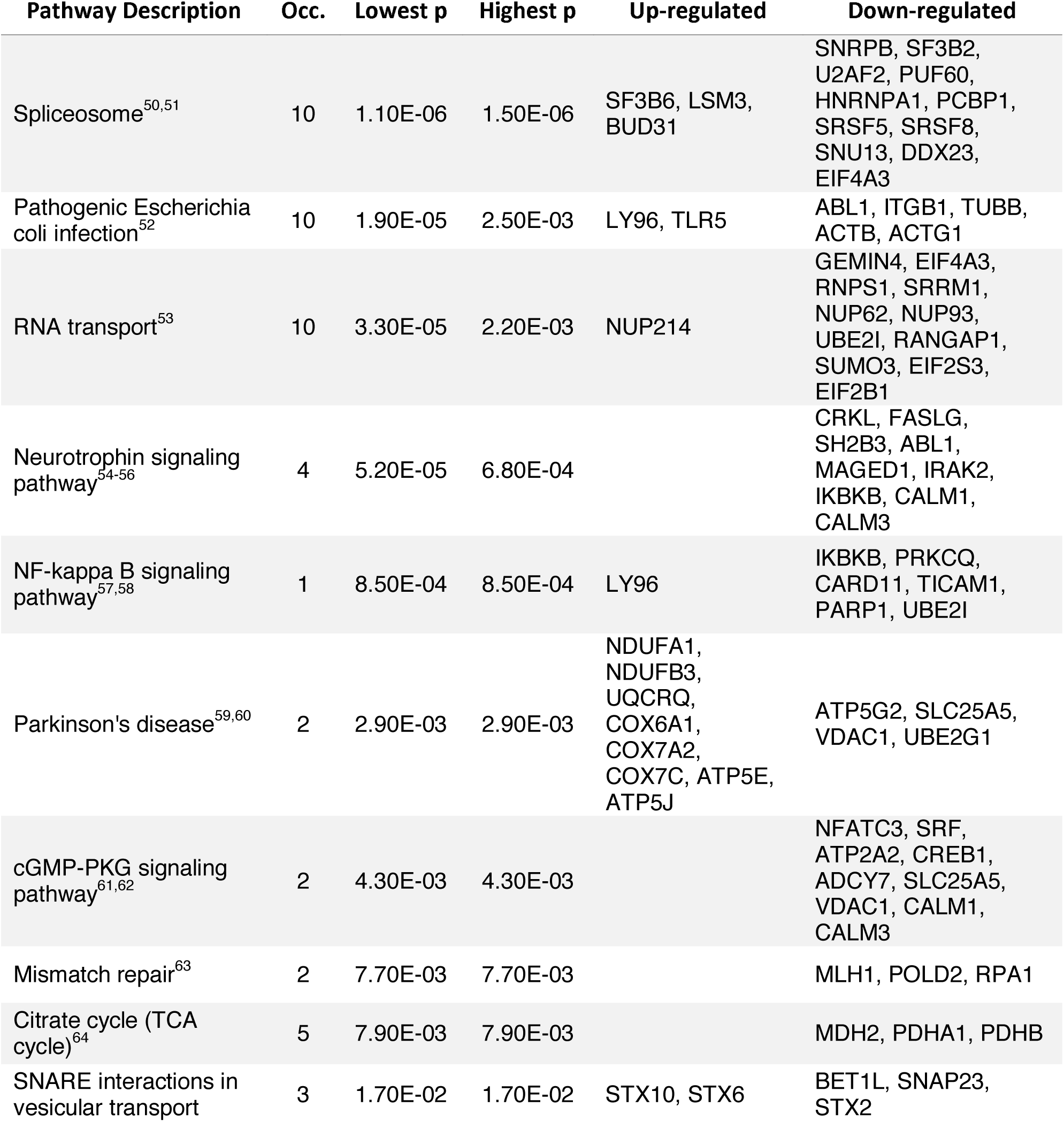
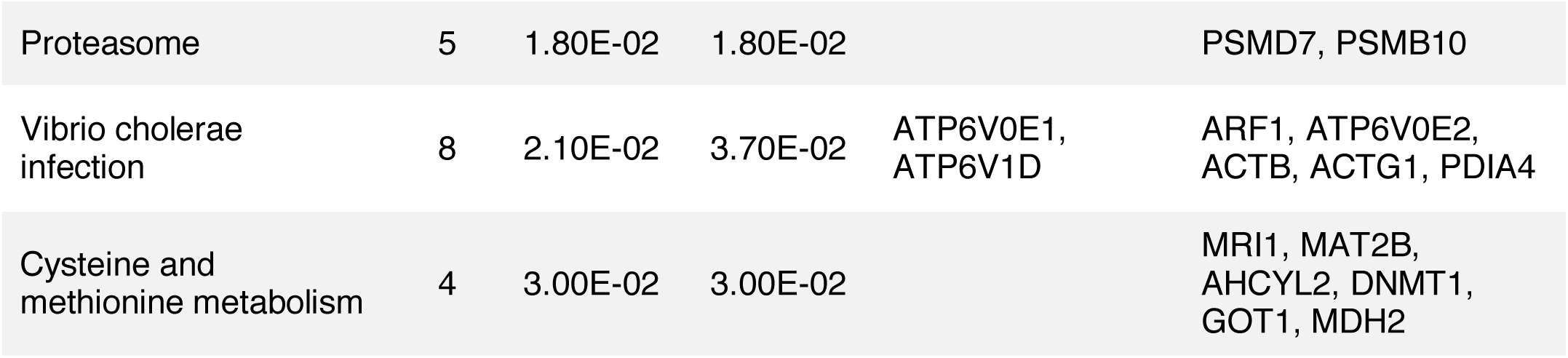
Representative pathways that were enriched in the RA differential-expression data. Pathway Description indicates the description of the given KEGG pathway. Occ. indicates the occurrence, i.e., the number of times the pathway was identified to be enriched over 10 iterations. Lowest p and Highest p indicate the lowest and highest p values calculated for the given pathway over all iterations. Up-regulated and Down-regulated indicate the up- and down-regulated DEGs that are involved in the given pathway.

**Figure 2:**
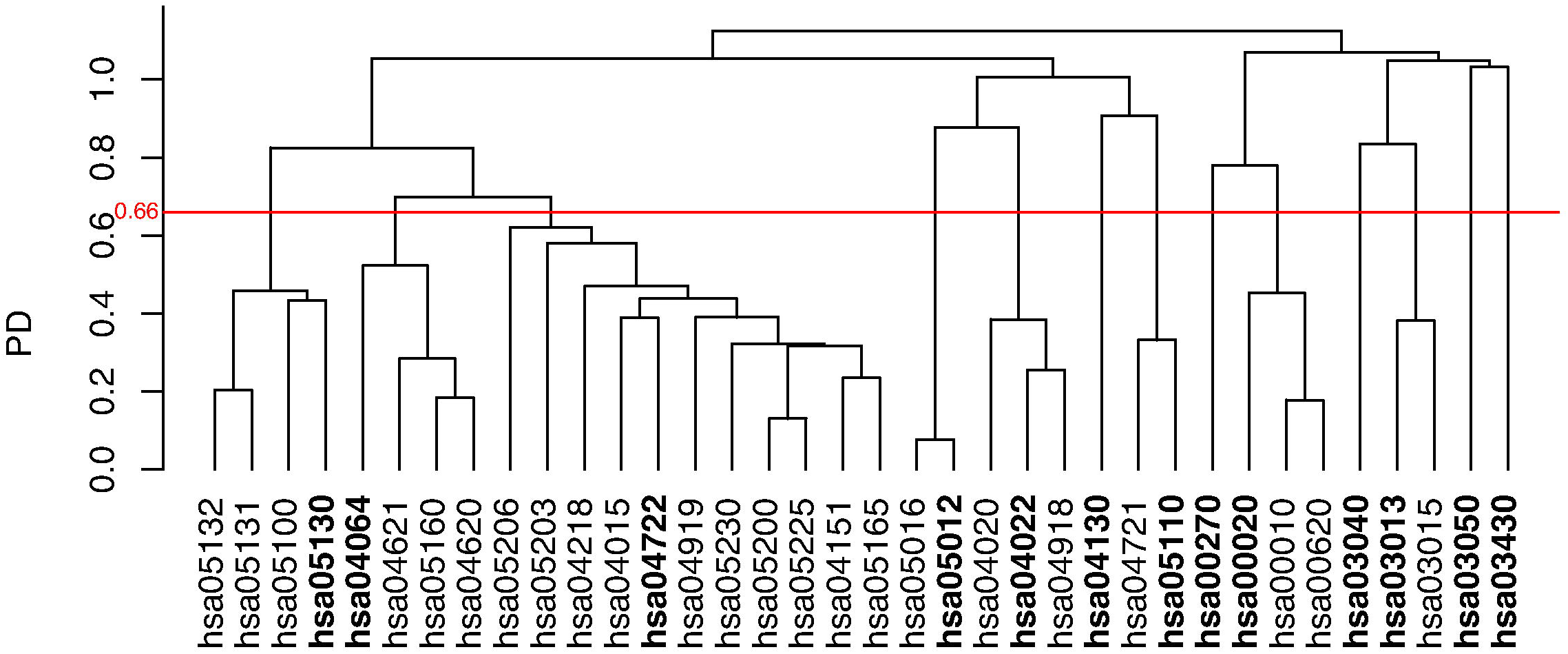
Clustering dendrogram of enriched pathways identified in the RA differential expression dataset. Vertical axis indicates the pairwise distance. The horizontal red line indicates the height at which the dendrogram is cut. The representative pathways, i.e. the pathways with the lowest p value in each cluster, are indicated as bold text.

The most significantly enriched pathway was “Spliceosome”. Autoimmune response to the spliceosome was previously reported in numerous autoimmune diseases, including RA^50^. Moreover, a recent study revealed there is a significant alteration of spliceosome components in RA patients^51^. This study suggested that alterations in the spliceosome could be associated with the development of RA and could also drive development of cardiovascular disease by altering the atherothrombotic profile in patients. “Pathogenic Escherichia coli infection” was found to be the representative pathway of the cluster, which also included the pathways, “Bacterial invasion of epithelial cells”, “Shigellosis” and “Salmonella infection”. The association between these pathways, suggesting a response to infection, and RA development is not certain. However, in a review on microbial infections and RA, it was reported that infections play an important role in the initiation and advancement of RA^52^. The review also discusses potential mechanisms whereby infection may promote the development of RA, such as generation of neo-autoantigens, molecular mimicry, and bystander activation of the immune system.

There is currently no study that explains the association of “RNA transport” with RA. However, a recent study that analyzed dysregulated genes in RA also found that DEGs were enriched in “RNA transport” among other pathways^53^. This implies that dysregulation of “RNA transport” may play an important role in RA.

The association between the “Neurotrophin signaling pathway” and RA is well supported by literature. A 2005 study compared nerve growth factor (NGF), brain derived neurotrophic factor (BDNF), neurotrophin 3 (NT-3), and neurotrophin 4 (NT-4) concentrations in the serum of spondyloarthritis (SpA), rheumatoid arthritis (RA) and osteoarthritis (OA) patients, and healthy subjects^54^. Significantly higher concentrations of NT-4 and lower concentrations of BDNF were reported in disease group compared to healthy controls. Another study investigated the mRNA expression of BDNF and NGF in synovial fluid cells of RA, SpA and OA patients^55^. It was detected that NGF was expressed at significantly higher levels in RA and SpA patients than in the OA group. A recent study that investigated the methylation patterns affecting the pathogenesis of RA identified that the differentially methylated genes participated in the “Neurotrophin signaling pathway”^56^ among others. This provides a further level of evidence on the involvement of this pathway in RA pathogenesis.

The “NF-kappa B signaling pathway” is known to play a key role in RA pathology. The transcription factor nuclear factor kappa B (NF-κB) is accepted as a pivotal regulator of inflammation in RA along with other aspects of RA pathology^57^. Studies in animal models of RA demonstrated the efficacy of inhibitors of this pathway. Therefore, the “NF-kappa B signaling pathway” is also considered a therapeutic target in RA^58^.

We identified the “Parkinson‘s disease” pathway as the representative pathway in the cluster which also included “Huntington’s disease”. The association between RA and neurodegenerative diseases is not entirely clear. However, a recent study investigated genome-wide pleiotropy between Parkinson‘s disease (PD) and autoimmune diseases and found a genetic link between PD and RA^59^. This study identified 4 loci with genetic risk variants conveying risk of both PD and RA. This genetic evidence supports our transcriptomic finding and suggests that PD and RA are affected by or induce similar biological processes. One of such common processes is likely inflammation: Known to be a key process in RA, inflammation is also reported to be etiologically involved in PD^60^.

“cGMP-PKG signaling pathway” enrichment is supported by two studies, which showed that RA shared epitope (an HLA-DRB1-encoded 5-amino acid sequence motif carried by most of RA patients) acted as a signal transduction ligand that interacted with cell surface calreticulin, triggered nitric oxide-mediated signaling events in opposite cells, and affected cGMP levels^61,62^.

The representative pathway “Mismatch repair” (MMR) consisted of only genes downregulated in RA. Supporting our finding, Lee et al. also identified suppressed MMR enzyme expression in RA^63^. This study also observed abundant microsatellite instability in RA synovium most likely due to MMR deficiency.

We also identified the “Citrate cycle (TCA cycle)” as one of the representative pathways. The cluster of this representative pathway also included “Pyruvate metabolism” and “Glycolysis – Gluconeogenesis”, suggesting a dysregulation in energy metabolism. This finding is supported by Yang et al. who screened different proteins and metabolites in the synovial fluid samples of 25 RA patients and 10 normal subjects to explore the pathogenesis of RA^64^. Ultimately, they identified energy metabolism disorder as a contributing factor of RA.

### 4. Conclusion

PathfindR is an R package that enables active subnetwork-oriented pathway analysis, complementing the gene-phenotype associations identified through differential expression/methylation analysis. Initially identifying active subnetworks in a list of significant genes and then performing pathway enrichment analysis of these active subnetworks makes the best use of interaction information between the genes. This, in turn, helps uncover novel in addition to known mechanisms underlying the disease, as demonstrated in the RA example.

As stated above, the pathfindR approach is based on PANOGA. This package extends the use of the active subnetwork-oriented pathway analysis approach to omics data.

Additionally, pathfindR provides numerous improvements and useful new features, listed in detail below.

To overcome inconsistent annotation issues, pathfindR converts gene symbols that are not in the PIN to alias symbols that are in the PIN. This ensures that the majority of genes from the experiment can be mapped to the PIN and the user can make the best use of the data at hand.

The package provides three active subnetwork search algorithms. The user is therefore able to choose between the different algorithms to obtain the optimal results.

For the greedy and simulated annealing active subnetwork search algorithms, the search and enrichment processes are executed several times. By summarizing results over the iterations and identifying consistently enriched pathways, the stochasticity of these algorithms is overcome. Because the genetic algorithm is time-exhaustive, it is executed only once.

In addition to the data frame object, the package provides an HTML report with links to a table of the active subnetwork-oriented pathway enrichment results and a table of converted gene symbols. The table of enrichment results contains links to the pathway diagrams of individual pathways. These diagrams display the involved genes colored by change values. pathfindR also allows for clustering of related pathways. This allows for further abstraction of the data and reduces the complexity of analysis.

All features in pathfindR work together to enable identification of dysregulated pathways that potentially reflect the underlying pathological mechanisms. We believe that this approach will allow researchers to better answer their research questions and discover novel mechanisms.

The pathfindR package is available on: https://github.com/egeulgen/pathfindR

---

**Supplementary Table 1:** Table of enriched pathways identified in the analysis of the RA differential-expression data with pathfindR. KEGG ID indicates the KEGG ID of the pathway. Pathway indicates the description of the pathway. Occ indicates the occurrence, i.e., the number of times the pathway was identified to be enriched over 10 iterations. Lowest p and Highest p indicate the lowest and highest p values calculated for the pathway over all iterations. Up_regulated and Down_regulated indicate the up- and down-regulated DEGs that are involved in the given pathway. Cluster indicates the cluster the pathway is assigned to upon clustering of the pathways. Status indicates whether the pathway is the representative pathway or a regular member in its cluster.

